# Controlling the synchronization and symmetry breaking of coupled bacterial pili on active biofilm carpets

**DOI:** 10.1101/2025.05.01.651758

**Authors:** Baha Altın, Enes Talha Günay, Yusuf Ilker Yaman, Alp Ünlü, Yiğithan Gediz, Neslihan Gedik, Bora Karadaş, Mustafa Başaran, Coşkun Kocabaş, Şahin Özdemir, Aşkın Kocabaş

**Affiliations:** Department of Physics, Koç University, Sarıyer, 34450 Istanbul, Türkiye; Sciences and Engineering Program, Harvard University, Cambridge 02138, MA; Department of Physics, Undergraduate Program, Boğaziçi University, 34342 Bebek, Istanbul Türkiye; Department of Materials, University of Manchester, Manchester, M13 9PL, UK; Department of Electrical and Computer Engineering, Saint Louis University, Saint Louis, Missouri 63103, USA; Koç University Surface Science and Technology Center, Koç University, Sarıyer, 34450 Istanbul, Türkiye

## Abstract

In the low Reynolds number regime, active biological systems utilize nonreciprocal cyclic activities to achieve motility, as seen in the spinning of bacterial flagella and the beating of cilia. Coupling among these active mechanical components leads to synchronization, and emergence of metachronal waves. Here, we report that biofilms of *Pseudomonas nitroreducens* form active carpet-like surfaces textured with diverse topological defects, generating Mexican-wave-like collective behavior in which bacteria periodically lift up. On these active surfaces, nonreciprocally coupled extension and retraction activities of bacterial pili drives these collective oscillations. Surprisingly, this collective behavior exhibits left-right asymmetry across the biofilm driving unidirectionally propagating waves. We discover that this directionality is primarily governed by an aging-related frequency gradient across the biofilm. Leveraging these insights, we further demonstrate the ability to control the collective dynamics of these waves, including symmetry breaking, transitions from spiral waves into target and propagating plane waves by manipulating the elastic properties of biofilms. Overall, our findings illuminate the fundamental role of nonreciprocally interacting active components in regulating synchronization, collective dynamics, and symmetry-breaking phenomena in biological systems.

## Introduction

At low Reynolds numbers regime, overcoming reversibility constrain to gain motility, is a challenging task which requires innovative solutions. Microorganisms utilize a variety of specialized active mechanical components to solve these difficulties by performing nonreciprocal motion^1–4^. Typical examples include the rotating helical flagella and beating elastic cilia. These active mechanical systems have the ability to execute cyclic motions, thereby breaking time-reversibility constraints^5^. In addition, due to the coupling among these active microstructures, complex synchronization phenomena also known as metachronal waves could emerge^6–8^. From these perspective coupling, synchronization and collective behaviors are intrinsically related and ubiquitous in nature. Such widespread dynamical processes are also biologically essential, as seen in cilia-driven mucus transport in lungs^9^, nodal flow in developing embryos^10^, central motor generators^11^ and spinal cord development^12^. These intricate synchronization behaviors share the same underlying dynamical principles that surprisingly align with non-Hermitian physics of active systems^13–16^. All these active systems are capable of injecting and transferring energy across the system by breaking certain symmetries.

Recent advances, especially those in the field of active matter physics^17–22^, have illuminated critical non-Hermitian features, such as chiral states where PT symmetry is broken. In this broken mode system could sustain the phase difference between fundamental fields or the modes which further trigger the directional energy transfer between the active compartments and generate travelling waves. Notably this non-Hermitian process is particularly driven by nonreciprocity in the coupled mechanical system^8,17,23^. Although non-Hermiticity has its roots in quantum mechanics^14,24^ and the fundamental principles have been broadly studied in photonics^25^, electronics, acoustics, optomechanics^26,27^, superconducting qubits^28^, trapped ions^29,30^, single-spin systems^31^, and in light-matter interactions^32^. This process is often achieved not only using strong nonlinearities, but also utilizing nonreciprocal interactions and coupling^33–35^. On the other hand, nonreciprocity is a very common feature in biological systems. Such as hydrodynamic interactions^36^ or prey predator^37^ relations could easily break action reaction symmetry and results in exotic collective behaviors. Extending the basic concepts from non-Hermitian physics to biological domain holds promise for shedding light on the fundamental principles of symmetry breaking in active and, more crucially, in complex biological systems.

Various biological models with specific active micromechanical components have been studied. The most notable biological platforms are ciliated epithelial cell^38^, bacterial carpet^39,40^, starfish embryo^41^ social amoeba^42–44^ and walking placozoa^45^. Addition to flagellum or cilia, bacteria could also use nano filaments known as type IV pili to gain motility on surfaces where the rotating flagella do not work effectively. Recent studies also highlighted the importance of pili controlling the collective behaviors of bacteria^46–49^ and also large-scale oscillations in the form of biofilm.

During our recent experiments, we noted striking dynamic patterns and unidirectionally propagating waves on the surface of *Pseudomonas Nitroreducens (PN)* bacterial biofilms. These unexpected collective activities ranging from spiral to planar wave formations, interestingly aligned with recent predictions of active matter physics^23,50^. These activities are reminiscent of metachronal waves observed in various biological systems. We also observed that these collective activities are driven by the coupled pili activity of the bacteria. Recently, large-scale pili-driven activities and spiral waves were also identified in *Pseudomonas aeruginosa*^49^. The main distinction of our observed metachronal wave lies in its localized behavior on the surface and broken left-right symmetry across the biofilm, which we refer to as the active biofilm carpet. On these active surfaces bacteria periodically lifting up and forming Maxican-wave like dynamics. In this study, we aim to provide a comprehensive experimental approach including biological, physical and theoretical tools to be able to understand and control these collective behaviors and also the symmetry breaking process of these biological systems.

Bacterial biofilms develop into multicellular communities characterized by interdependent biological structures. Understanding the dynamics of these dense biological formations is vital, given that their collective actions can amplify their pathogenic potential. The collective movement of pili and its variability among strains offers a crucial foundation for understanding the influence of pili activities on pathogenicity. Moreover, pili is also a primary target protein for next-generation medications and antibacterial therapies. In essence, comprehending the effect of pili dynamics on the collective behavior of biofilms can greatly influence human health by preventing severe infections.

## Results

### Emergence of metachronal waves on active biofilm carpets

Bacterial biofilms are commonly used in various experiments particularly to investigate host-pathogen interactions. During our recent screen of biofilm library for nematode *C.elegans*^51^, we observed that *Pseudomonas Nitroreducens* (PN, Materials and Methods) bacterial biofilms exhibit rhythmic activities approximately 10 hours post-inoculation (Figure1a-b). In their firing phase, faint propagating pairs of spiral wave patterns emerged on the biofilm’s surface. These waves have a periodicity of around 2-7 minutes (Figure1c). We also determined that the visibility of these waves improves dramatically with oblique contrast and polarized light-based microscopy (Figure1a-b, Supplementary Video1-2). Later on, these waves were very dynamic and converged to various combinations of spiral, target and planar waves. The most striking feature of these waves is their broken left-right symmetries. They, often unidirectionally propagate from right to left (edge to the center) or radially in inward direction across the colony (Figure1a-b). Furthermore, the contrast of the wave’s switches between the dark to bright sharp lines depending on the direction of propagation. This suggest that the front kink of these waves scatters the light asymmetrically (Supplementary Figure 1). We observed that these waves predominantly propagate on the biofilm’s surface rather than its bulk region (Figure1e). This critical feature is particularly revealed by the nature of our optical imaging systems which is very sensitive to the surface topography. To validate this observation, we employed cytoplasmic GFP for imaging. Yet, no waves were detected in the green fluorescence channel (Supplementary Figure2). Similarly, when we labeled extracellular DNA with a dye, the waves remained unobserved in the fluorescence signal (Supplementary Figure2). Moreover, close-up imaging (100X) highlighted the bacterial displacement together with clear optical dark or bright contrast on the surface (Supplementary Video3). Imaging growing biofilm starting from a single bacterium clarifies that oscillations first starts from dense regions and gradually propagate through the entire colony (Supplementary Video4). Further motivated by recent studies^48,49^, we also tested the *Pseudomonas aeruginosa* strain PA14 under the same conditions. Similarly, PA14 exhibited weak but comparable wave patterns, though within a shorter time window (Supplementary Figure4).

**Figure 1.**
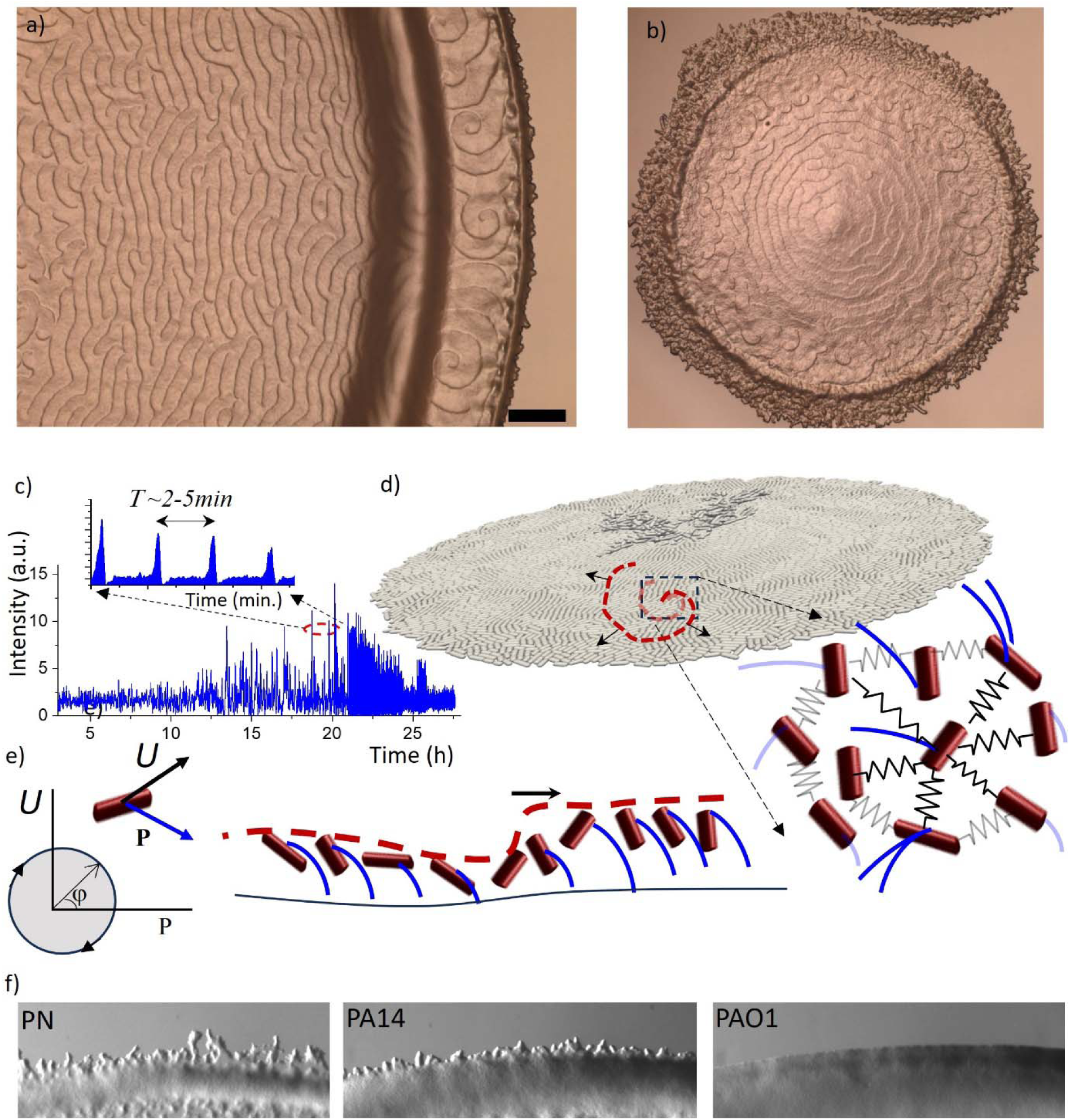
Emergence of unidirectionally propagating waves on bacterial biofilms. (a-b) Optical imaging shows spiral, planar (a), and radially shrinking waves (b) propagating unidirectionally on biofilm surfaces. Left-right or radial symmetry is generally broken on growing biofilm surfaces. Scale bar 250 um. (c) Time response of optical scattering signal indicating the firing state of coupled pili dynamics. (d) Schematic representation of the coupling mechanism of pili on biofilm surfaces, modeled as an active solid. Red cylinders represent the bacteria in the biofilm. The oscillatory extension and retraction of pili act as active units. (e) Schematic representation of the collective behavior of elastically coupled active biofilm surfaces, characterized by local displacement (//) and pili polarization (P) undergoing limit-cycle oscillations. Propagating waves remain localized on the surface and travel toward the direction of the sharp rising edge. f) Optical images of the leading edge of the growing biofilm with fingering instabilities. PN and PA14 show strong fingers compared to PAO1.

To delve deeper into these phenomena, our next focus was on the necessary media, given that the original experiments were conducted on *C. elegans* using Nematode Growth Media (NGM) plates (see Materials and Methods). Interestingly, we found that these activities were exclusive to NGM. Substituting NGM with a standard LB plate did not reproduce the same dynamics; instead, the bacteria grow rapidly. Yet, upon omitting yeast extract and replacing bactotryptone with bactopeptone, a major nutritional component, from the LB media PN resumed its activity (Supplementary Figure6). These findings underscore the significance of nutrient restriction as a primary factor. Remarkably, these observations are unexpected, but become evident due the use of NGM plate, which is not common in microbiology experiments.

Although PA14 generates similar waves we observed that PAO1, a close relative of PA14, failed to produce waves on both NGM and LB plates (Supplementary Figure5). Worth noting, PAO1 is also an important human pathogen causing *Cystic Fibrosis* (CF). Finally, through this comprehensive screening effort, we successfully identified a group of bacteria, offering the potential to examine various factors, from genetics to physical domain, influencing the emergence of these spiral waves. Both PAO1 and PA14 possess robust genetic toolkits, while PN strain display a clear and prolonged active carpet state for detailed physical investigations. The other critical difference between all these biofilms is the finger formation around the leading edge. Unlike PAO1, PN and PA14 form very strong instabilities leading unstable extensions (Figure1f).

To gain deeper insight into the mechanisms underlying wave formation, we imaged the dynamics of individual bacteria from the fingering regions toward the center of the biofilm. This distinction is critical because, unlike the biofilm center, the edges do not generate waves. We observed that bacteria near the fingering regions remain motile and exhibit collective flow. In contrast, bacteria at the biofilm center are surface-attached and undergo periodic lifting motions. This behavior strongly resembles Mexican-wave dynamics (Supplementary Video5, 6).

We further found that the central region of the biofilm is mechanically more elastic (Supplementary Figure3), whereas the edge regions—where wave formation is absent—are motile. These observations suggest that gradual biofilm maturation is a key factor that transforms motile bacteria into a periodically moving but spatially constrained state. Consistent with this picture, the PAO1 strain, which has a strong biofilm-forming capability, completely suppresses surface oscillations. In contrast, the PA14 strain exhibits intermediate behavior, sustaining a partial transition between motile and locally constrained dynamics. Remarkably, signatures of this transition and wave generation are already detectable at the earliest stages of finger formation.

We then shifted our attention to the genetic distinctions between PN, PA14 and PAO1. Initially, we first explored the contribution of bacterial pili and flagella as an active mechanical structure. We found that PA14:ΔPilB and PA14:ΔPilA mutations eliminated the waves but not PA14:ΔFilK (Supplementary Figure4). This suggests that activity is exclusively originating from pili dynamics and the original dynamics is similar to the recent work^49^, but activities are localized to the surface to the transition from motile to biofilm forming state. We then explored the potential of hyper piliation, which might amplify the coupling of pili on the biofilm surfaces. Yet, the PAO1 ΔPilH mutant, known for its polarized and hyper piliation condition^46,47^, did not manifest the waves but provided only small fluctuations on the surface (Supplementary Figure5). Then, we pinpointed that a significant variation was in the pili subunit groups^52^. While PA14 possesses group 3 Pilin, PAO1 has group 2 Pilin. It’s essential to highlight that all these bacteria have type IV pili, though they might be categorized into separate groups. Currently, our hypothesis posits that diverse pili types might exhibit varied physical characteristics such as elasticity or rigidity. Our understanding of PN at the genetic level remains limited but it shares strong similarity with PA. Nonetheless, the various genome sequences of PN are accessible. Our analysis of their pili related genomic region revealed a resemblance to the group 4 and 5 pili (Supplementary Figure7). All these firing pili groups 3 and 4, 5 have additional accessory genes in the genome and they may contribute to the expression or folding of these pilins. Together with the unstable edge fingering (Figure1f), the comparison of PN and PA strains support the idea that group of pili and particularly pili expression play a critical role in driving waves on these bacterial biofilms.

### Modelling of propagating spiral waves

To get more intuition about the dynamics of the waves and emergence of the broken symmetry across the biofilm, we next focused on the mathematical modelling. From an active matter perspective, spiral waves can emerge in non-equilibrium and excitable media^53–55^ as well as in coupled oscillatory systems with delays^56^. Despite these systems exhibiting similar wave patterns, the internal mechanisms that trigger the waves vary significantly. Recent studies particularly on the concept of nonreciprocity provides comprehensive approach^23,50^. It is important to note that coupling between active mechanical units could introduce nonreciprocity into the system due to local symmetry-breaking processes. Inspired from the previous studies on the dynamics of cilia^17,57^, and recent robotic system^58^ we proposed that coupled active pili system could also show similar limit cycles oscillations defined by bacterial displacement (**U**) and the polarization of the bacterial pili (**P**) (Figure1e). We note that combining phenomenological field models with more detailed elastic solid models could provide a more comprehensive framework for understanding how the elastic properties of the biofilm drive these collective oscillations. To study these phenomena, we first utilized a minimal phenomenological phase field model that includes a nonreciprocal coupling term. The oscillatory dynamics of the model are driven by oscillatory extension and retraction processes. The Kuramoto based model^23,56,59–61^, which effectively captures the phase dynamics of coupled oscillators, is extended here by incorporating a delay term (α) that disrupts the odd symmetry—representing action and reaction symmetry between interacting oscillators—and enriches the system’s dynamic repertoire. This modified model is described by the equation:

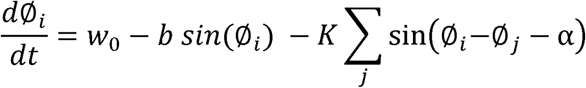

Here, (ø represents the local phase of individual pili oscillations (The phase of confined parameter space defined by **U** and **P**, Figure 1e). We assume interactions only among nearest neighbors, where W_Q_ represents the uniform intrinsic oscillation frequency of an isolated oscillator. The term α specifically break odd interaction symmetry between pili, facilitating the formation of travelling waves. Note that traveling waves indicate this broken PT symmetry between these fields. It’s important to emphasize that the detailed biophysical mechanisms behind this nonreciprocity remain unclear. Several possibilities, such as force relaxation dynamics or hydrodynamics interactions, could also result in similar nonreciprocal behavior^62^. Our phenomenological model accurately captures the principal features of these phenomena. While reciprocally coupled oscillators (α = 0) tend to achieve a globally synchronized state (Figure 2a), nonreciprocity favors the formation of traveling waves (Figure 2b-c, Supplementary Video7, Supplementary Figure8). The excitability term (**b**) shapes the pulsative nature of the waves (illustrated in Figure 2d-e) by amplifying the effects of specific phases within the coupling mechanism^56,63,64^.

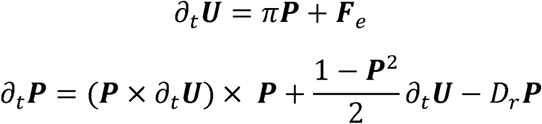

**Figure 2.**
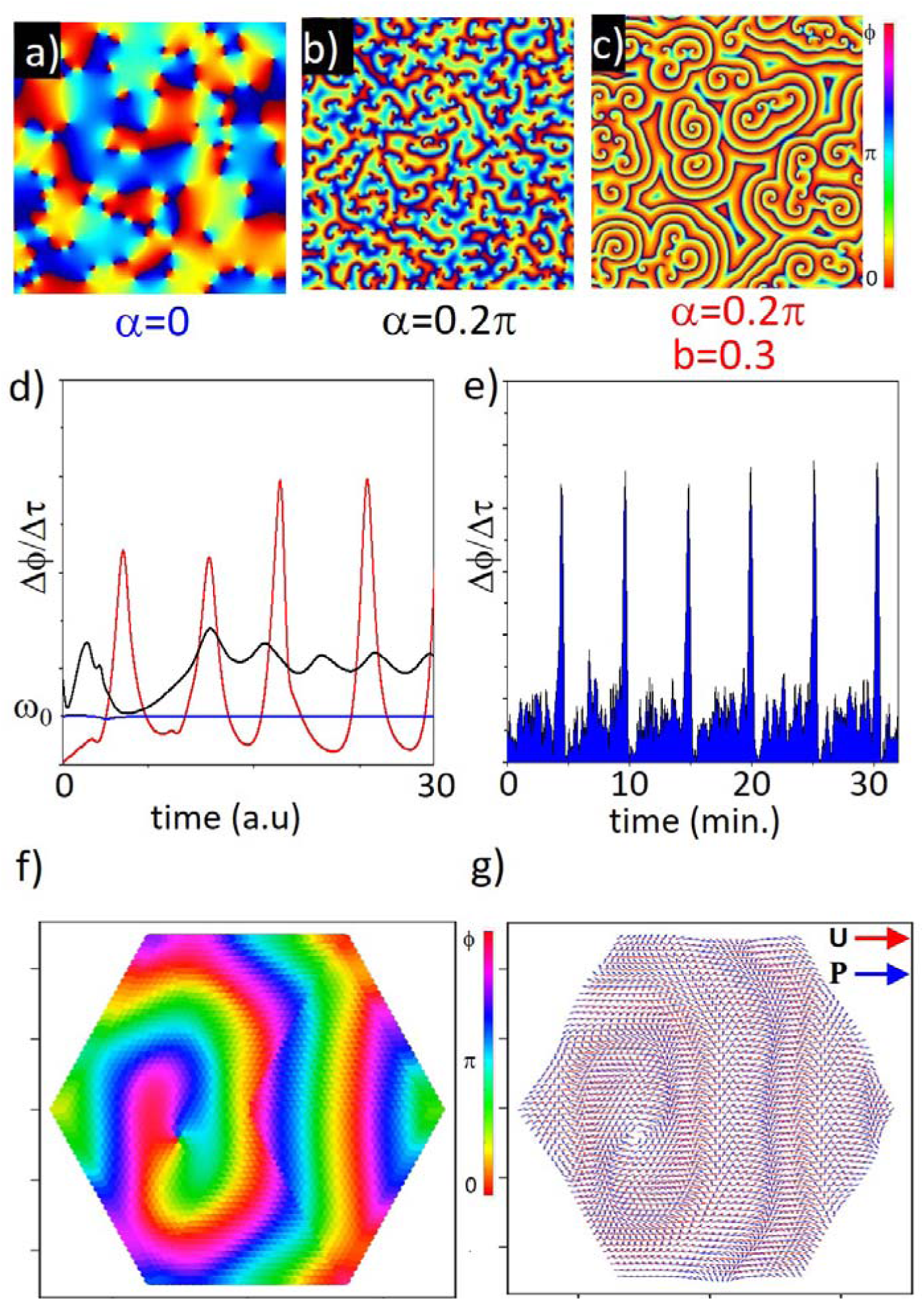
Numerical modeling of coupled pili dynamics as an active carpet. (a-c) Numerical simulation results based on the nonreciprocal Kuramoto phase-field model. Nonreciprocal coupling term (_α_) among bacteria drives the emergence of spiral waves, while the excitability term (b) of the mechanical system induces pulsatile behavior. (d) Numerical simulations of local oscillation frequencies under varying conditions (_α_ = 0, _α_ = 0.2_π_, and b = 0.1_π_). (e) Experimental measurements of pulses on biofilm surfaces confirming pulsatile responses. (f-g) Numerical simulations of elastically coupled biofilm structures using the active solid model. Active solids exhibit large-scale spiral and planar wave formations. Displacement (U) and pili polarization (P) fields highlight the essential phase difference necessary for wave propagation and limit-cycle oscillations.

Next, we turned our focus to continuum simulations to capture the essential properties of these dynamic processes (Supplementary Video8). In our previous experiments, we observed that changes in the elastic properties of the colony during the transition from a motile state to a biofilm-forming state are critical. The recently developed active solid model provides a broad and robust framework for studying collective behaviors of interacting elasto-active components, spanning from robotic systems to human crowds^58,65^. The core idea of this model is based on two nonreciprocally coupled fields: displacement (**U**), representing bacterial displacement within the biofilm lattice, and polarization (**P**), representing the orientation of pili, which exert active forces whose direction can be modulated by the displacement field (Figure1d). The dynamics of the active biofilm lattice can be described by coupled equations, where **F_e_** denotes elastic force and D_r_ represents polarization relaxation. The underlying rationale of these equations is that retracted pili exert active forces on bacteria, which are elastically coupled to the biofilm structure. Local biofilm displacement can guide the growth direction of pili extension, influenced by local deformation. Importantly, since pili are anchored to the substrate, their orientation dynamically adjusts based on extension processes. Our simulations demonstrate that the active solid model successfully captures critical dynamical phenomena, such as the formation of spiral and planar waves (Figure 2 f-g). As anticipated, fields **U** and **P** develop a characteristic phase difference, indicative of limit cycle oscillations. Moreover, analysis of vector configurations around defect cores reveals directional reversals of displacement **U** on opposite sides of defects, highlighting the fundamental mechanism underlying defect formation, where **U** diminishes, and the phase (ø) becomes undefined (Supplementary Figure9). We should note that traveling waves indicate broken PT symmetry between these fields triggered by nonreciprocity, with spiral waves serving as a classic signature of this phenomenon. A further transition from spiral to planar waves reflects an overall asymmetry in the frequency profile, which is not directly related to PT-symmetry breaking.

### Controlling the transition between the waveforms on active biofilm carpet

Extensive parameter testing has revealed that our models not only capture the emergence of spiral waves but also predicts transitions into target and plane wave solutions which we commonly observe in grooving biofilms. This transition is particularly emerge around the edge of the plate where the drying process is gradually modifying the agar plate. During the transition to target waves, two oppositely spinning spiral waves with topological charges of +1 and -1 merge, forming a symmetrically expanding target wave. Subsequently, fast oscillating plane waves progressively dominate the entire simulation domain (Figure3 a-c, Supplementary Video7).

Motivated by our simulation results and experimental observations, we hypothesize that our active carpet system could also facilitate similar transitions between different forms of traveling waves^66,67^, offering a means to externally control the dynamics of the system and may explain the details of broken symmetries. We observed that adding a water droplet to the biofilm surface led to the transformation of spiral waves into target and planar waves (Figure 3d-f). To recover the spiral waves, we increased the surface temperature by gently heating the system. As expected, with the temperature rise, the leading edge of the planar waves first became noisy, and multiple target waves randomly appeared then finally converge to spiral waves (Supplementary Video9). These findings are in interesting similarity with recent theoretical predictions in different nonreciprocal systems^50,66^.

**Figure 3.**
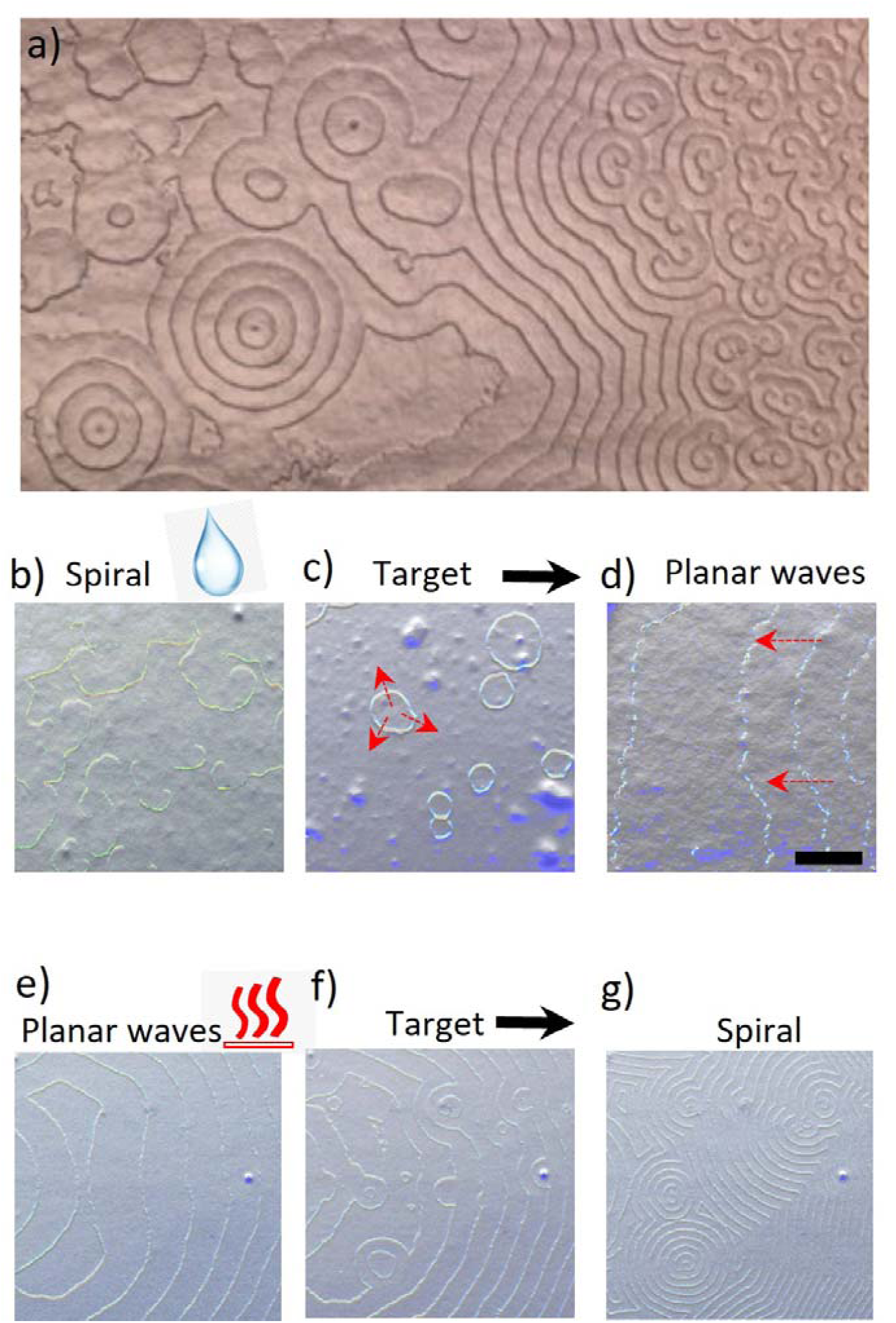
Controlling transitions between spiral, target, and planar waves. (a) Experimentally observed transitions from spiral waves to target and planar waves spontaneously emerge across the plate. Pairs of spiral waves merge, forming topologically neutral target waves, which eventually give rise to planar waves dominating the biofilm surface. (b-c) Optical imaging of biofilm surfaces demonstrating similar controlled transitions between spiral, target, and planar waves experimentally triggered by adding a water droplet. Red arrows indicate wave propagation direction. (e-g) Controlled recovery of spiral waves achieved by heating biofilm surfaces, removing excess moisture, and facilitating re-emergence of spiral waves around a specific inhomogeneities.

Additionally, by incorporating the water-soluble polymer polyethylene glycol (PEG) into the biofilm, we managed to not only alter wave propagation patterns but also direct the movement of inward growing target waves. Intriguingly, the addition of small drop of PEG modified the intrinsic oscillation period of the pili dynamics, creating a radial gradient profile largely due to the slow evaporation rate and uneven deposition of PEG on the surface. This inward wave guidance is particularly controlled by frequency gradient (Figure 4a-d). The center has slow oscillations than the edge. This gradient profile is the critical feature to understand the basics of broken symmetry observed in the circular biofilms. In naturally growing colony oscillation frequency is also varying (Figure 4a-b). We hypnotize that this variation is age dependent where the center of the colony is slowing down due to change in elastic properties of the biofilm. Before delving deeper into this interesting symmetry-breaking process, we investigated how the aging of the biofilm affected the oscillation dynamics. This is because symmetry breaking occurs at late stage where the aging is critical. To do so, we first measured the oscillation periods as the uniform colony aged. We observed that the periods of these spiral waves continuously increased (Figure 5a-b). These results suggest that the intrinsic oscillation dynamics of the Pili decreased, likely due to biofilm formation which changes the elastic properties of the colony, during first 24 hours.

**Figure 4.**
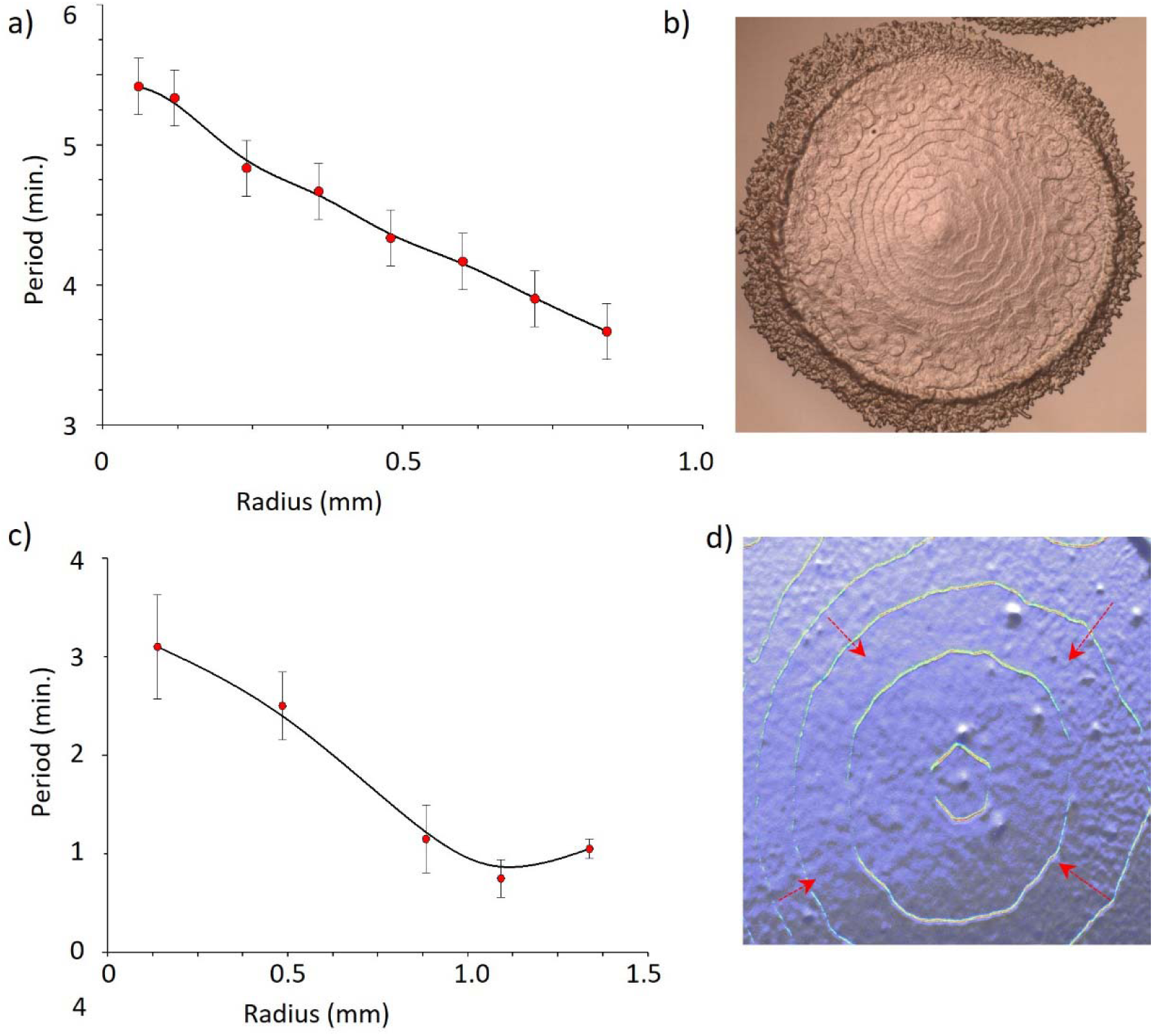
Controlling dynamics of inward propagating waves. (a) Optical imaging of inward propagating waves within a circular biofilm structure. (b) Period of pili oscillations decreases toward the colony center. (c) Application of a small droplet containing PEG creates a radially varying period profile, guiding inward wave propagation, capturing similar wave the dynamics on naturally growing radially symmetric biofilms. Error bar shows the S.D, N= 5 measurements.

**Figure 5:**
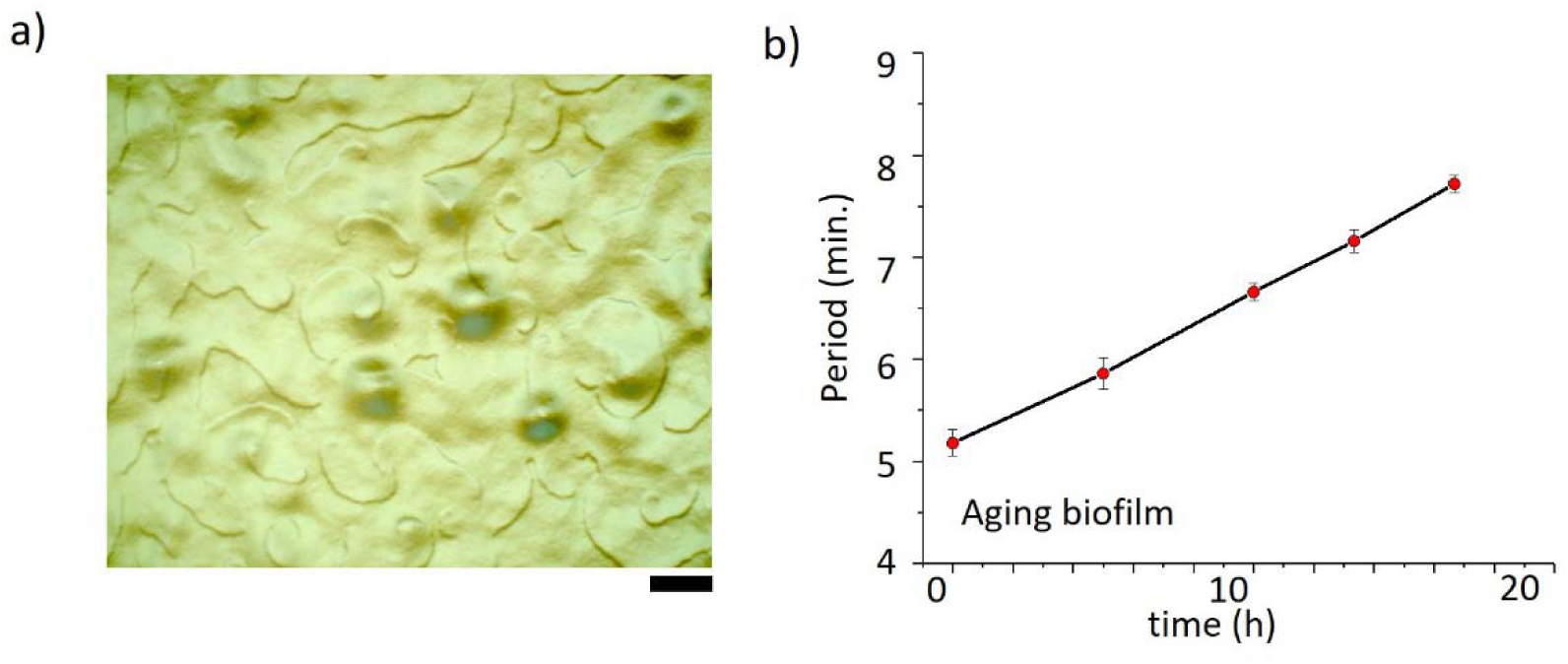
Age-Dependent Dynamics of Oscillations. a) Sample image showing multiple pairs of spiral waves on a uniform biofilm surface. Scale bar: 100 _μ_m. b) Period of the oscillations increases as the biofilm ages. Error bar shows the S.D. N= 10 measurements.

Further, we examined the growing biofilm starting from a thin inoculation strip, we reproducibly achieved a broken left-right symmetry for plane waves across the colony (Supplementary Video10). The waves propagated from the edges toward the center. This asymmetry became particularly pronounced a few days after inoculation, during which the colony edges continued to grow while the center remained stationary (Figure 6a, b). To finally confirm this observation, we numerically solve the coupled oscillators under varying frequency. Similarly, we found similar left-right asymmetry where the planar waves propagating towards the slower region of the biofilm center (Figure 6c, d, Supplementary Video11).

**Figure 6.**
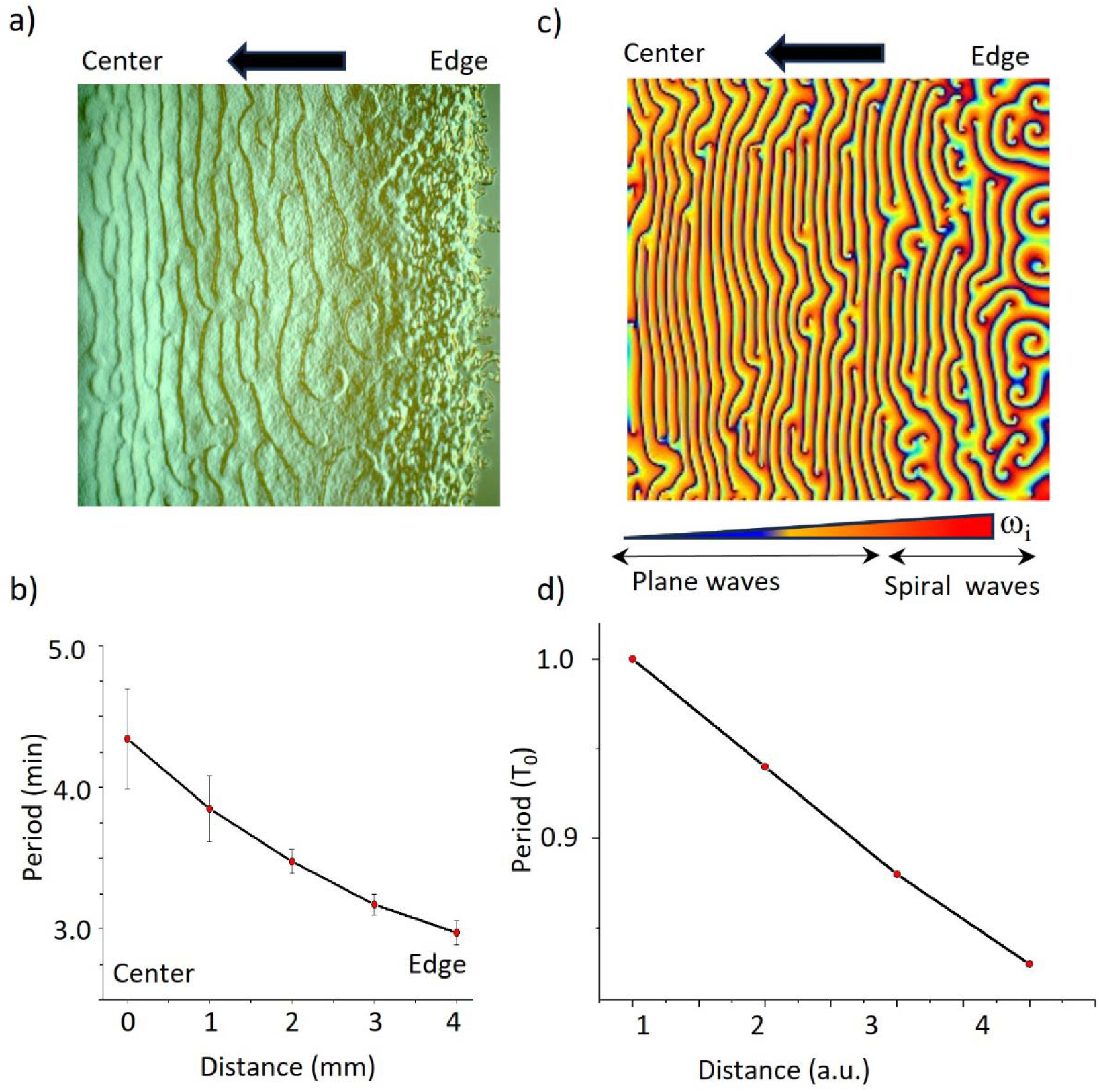
Left-right asymmetry in naturally growing biofilms. (a) Representative image of a bacterial biofilm at a late growth stage (3 days post-inoculation). Metachronal waves propagate towards the biofilm center, with the leading edge displaying chaotic dynamics. Following growth cessation, spiral waves gradually converge into planar waves propagating inward. (b) Oscillation period increases towards the biofilm center. Error bar indicates the S.D. N= 10 measurements (c-d) Numerical simulation demonstrating the formation of planar waves and the left-right symmetry-breaking process, driven by a spatially varying intrinsic oscillation frequency(d), capturing dynamics observed in naturally growing biofilms.

### Defect dynamics controlling the transition between spiral to target waves

To better understand the dynamics of the transition between different form of the waves we focused on numerical simulations. We noticed that the motility of defects is the crucial parameter governing the transition between spiral, target, and planar waves varying the moisture content provides an effective and experimentally accessible control this motility. Our analyses revealed that spiral defect cores can move and merge to form target waves or annihilate entirely—processes that we also observe experimentally. This rich dynamical behavior underscores the importance of elasticity in shaping pattern transitions. First, we compare defect dynamics in both Kuramoto-based simulations and the active solid model. Both systems exhibit similar defect-survival behavior. As shown in Supplementary Figure10, pairs of unlike (+/−) defects can stably persist only at high nonreciprocity. We further quantify this behavior by plotting the separation distances between unlike defect pairs and find that short-range defect separations are possible exclusively in the high-nonreciprocity regime (Supplementary Figure11). This high-nonreciprocity regime corresponds to the dry biofilm state. Increasing moisture reduces elasticity, leading to the loss of stable defect dynamics and promoting the annihilation of unlike defect pairs, which in turn drives the system toward target-wave formation and ultimately planar waves. Conversely, heating the biofilm removes water, enhances elasticity, and increases the system’s ability to sustain closely separated defect pairs. Experimentally, we further observe that removing water by heating enhances surface nonuniformities, which readily trigger defect-pair formation (Supplementary Video9). To investigate this mechanism, we performed additional simulations in which local nonuniformities were introduced (Supplementary Video12-13). Consistent with experiments, defect-pair generation occurs only at high nonreciprocity, where pairs of unlike defects can be stably maintained. Experimental observation (Supplementary Video9) also show that surface nonuniformities on the biofilm surface similarly trigger the formation of closely separated defect pairs.

### Controlling left-right asymmetry

As a final step we focused on how to control the symmetry breaking and dynamics of metachronal waves. We noticed that temperature adjustments also reversibly effect the oscillatory behavior of the waves which increased the oscillation frequency (Figure 7a, b). This suggest that temperature could be also used as a complementary technique to dynamically guide the wave propagation by controlling the frequency gradients. To do so we embedded metallic pipes in the bulk agar plate and tuned the local temperatures (Supplementary Figure13). Setting different temperatures in these pipes creates temperature gradient (∇T∼10°C/cm). This gradient profile converted spiral waves quickly to planar waves, propagating from warmer (31°C) to cooler (21°C) areas. Similarly, when we switched the temperature profiles wave propagation also reversed (Figure 7c-d). This result indicates that spatial temperature variations can effectively dictate wave propagation directions. These comprehensive observations confirm that the developed frequency gradients of intrinsic Pili oscillations is the critical feature breaking symmetry and controlling the propagation directions of the waves.

**Figure 7:**
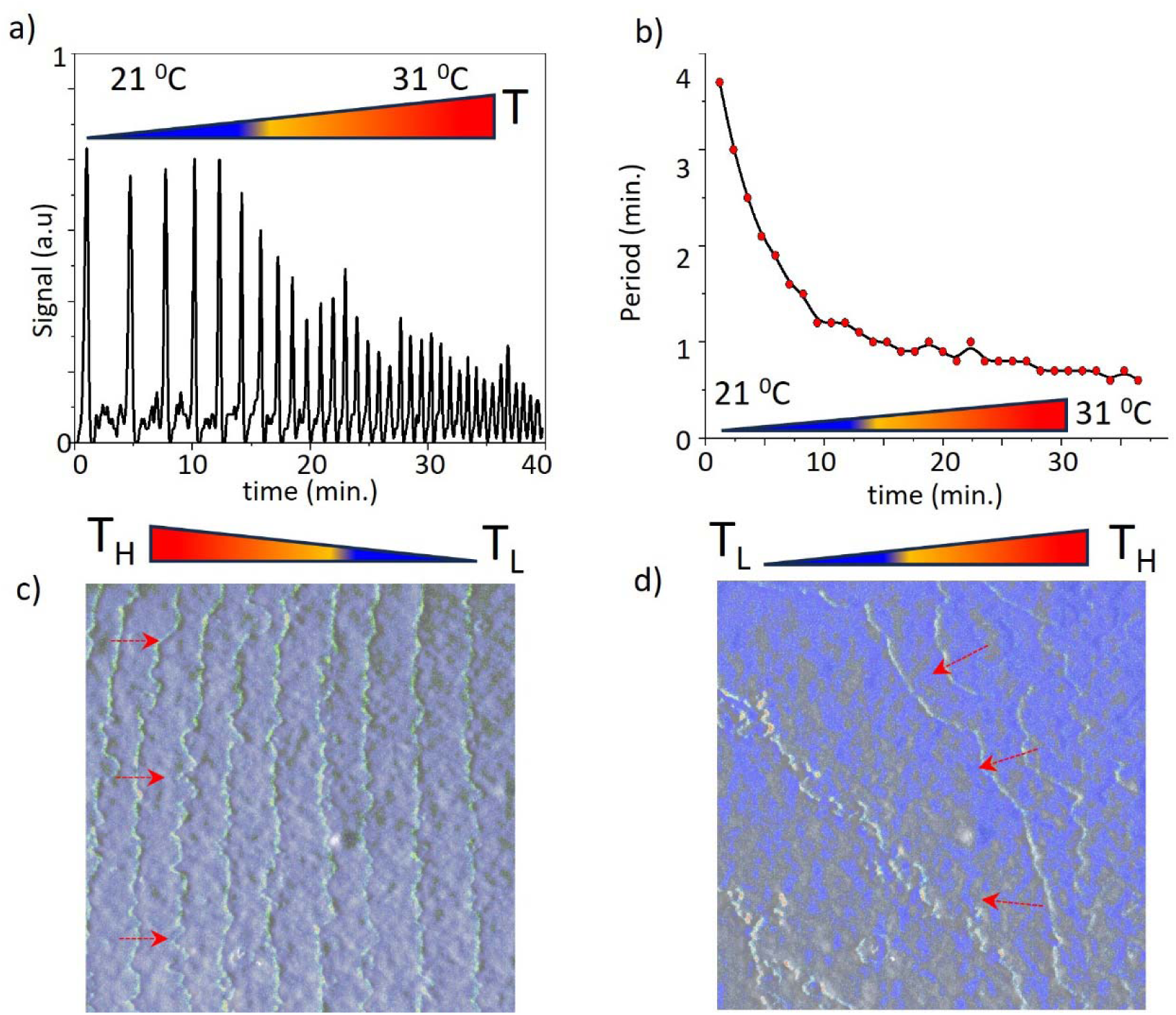
Temperature-Controlled Dynamics of Metachronal Waves generating left-right asymmetry. a) Optical pulses from spiral waves on the biofilm surface as temperature increases from 21°C (T_L_) to 31°C (T_H_). b) An increase in temperature raises the oscillation frequency c, d) Controlling the propagation of the waves by creating a temperature gradient. Waves propagate from the warmer area (fast oscillating, T_H_) to the colder region (slow oscillating, T_L_). d) Reversing the temperature gradient changes the direction of propagations of the waves.

## Discussion

Synchronization is a ubiquitous phenomenon in biological systems. A classic hallmark signature of this dynamical process is global synchronization, where all components of the system tend to merge in phase, as observed in fireflies, clocks and metronomes. Hydrodynamics or intrinsic activities of the interactions, however, introduce nonreciprocity into these tightly coupled systems, fundamentally altering the synchronization process. This action reaction asymmetry drives the emergence of large-scale propagating waves, often emerge as spiral waves in two dimensions. Remarkably, diverse systems from proteins in cell membrane to social amoebae exhibit similar dynamics, highlighting the special importance of these waves in biological systems particularly due to their transport capabilities.

From a symmetry perspective, nonreciprocity is the key feature in understanding these phenomena. Recent advancements in active matter physics have bridged various seemingly unrelated fields around this concept. Particularly, non-Hermitian physics, including PT-symmetry, provides a beneficial macroscopic metric for understanding a system’s response through the classical formalism borrowed from quantum physics and optics. Despite the diverse nature of the dynamic processes and the varying types of interactions—from predator-prey dynamics, specific chemical reactions, to elasticity of active metamaterials—the common feature across these systems is to maintain phase difference between the fundamental fields that support limit cycle oscillations enabling collective motility and transport capabilities to the interacting active system.

We have observed that bacterial biofilm surfaces also known as active carpet exhibit similar dynamics due to coupled and cyclic pili extension and retraction activities. Although these nanoscale activities make the biophysical mechanisms complex and demanding to elucidate, the fundamental characteristic of the process remains consistent: the cyclic motion of bacterial pili forms limit cycle oscillations in phase space to disrupt local or action-reaction symmetry among the interacting components.

Using a phenomenological model, we have concentrated on identifying critical parameters that control the system dynamics and synchronization. We have also developed methods to manipulate and control the collective dynamics and propagation of metachronal waves by adjusting these critical parameters. From the perspective of active matter, the controllability of densely coupled active mechanical components is crucial, especially at low Reynolds numbers where controlling turbulence and managing long-range transport is vital. Biological systems can effectively address these challenges and precisely control active and programable transport^68^. A deeper understanding of the underlying principles and critical symmetries governing this controllability could provide new insights into the complexities of biological systems. We speculate that these waves could promote active diffusion of oxygen and DNA. Further biophysical investigations are needed to delve deeper into these fascinating collective behaviors and its biological significance. Furthermore, the dynamical similarities between non-Hermitian physics and PT symmetry and the metachronal waves due to nonreciprocity are remarkable. Finally, we also hypothesize that the concepts developed in no-Hermitian physics could bring new perspective to dissect the complexities of these collective behaviors. Finally, our findings also raise several questions, particularly regarding the biological significance of these waves in the physiology of the biofilm. Rhythmic activities^69,70^ play a critical role in bacterial communities. We hypothesize that this synchronization may facilitate rhythmic behavior across the biofilm or enable expression and the transport of specific biomolecules under stressful conditions. Further studies are needed to elucidate the mechanisms underlying this emerging phenomenon.

---

**Supplementary Figure 1.**
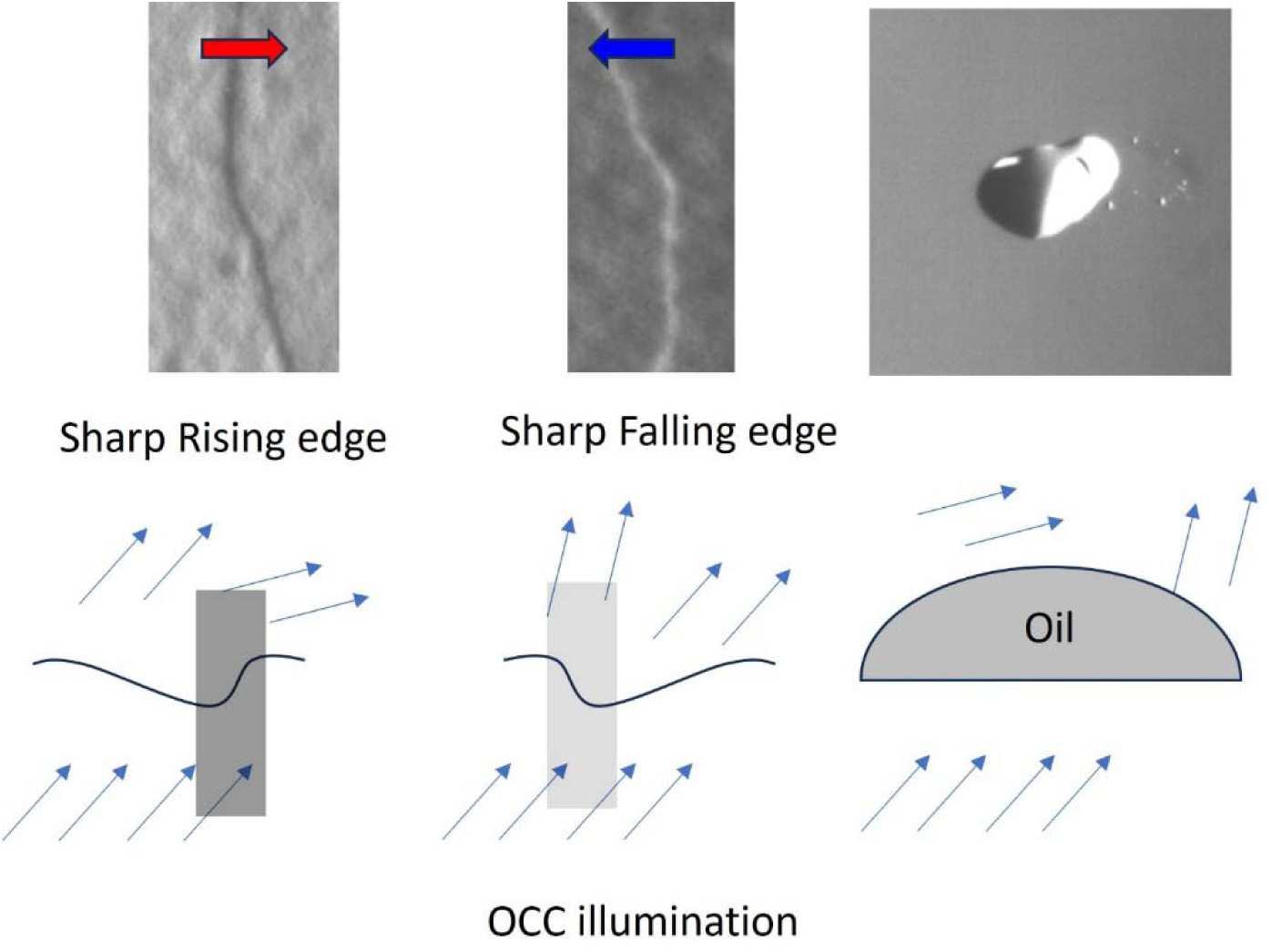
Asymmetric surface deformation drives directional propagation. OCC imaging provides clear dark and bright optical contrasts, originating from oblique illumination and asymmetric scattering by surface waves. An oil droplet was used to investigate the shape-dependent propagation mechanism. Under oblique illumination, sharp rising edges generate dark contrasts, while falling edges create bright contrasts. Sharp edges correspond to rapid pili retraction, followed by gradual recovery.

**Supplementary Figure 2.**
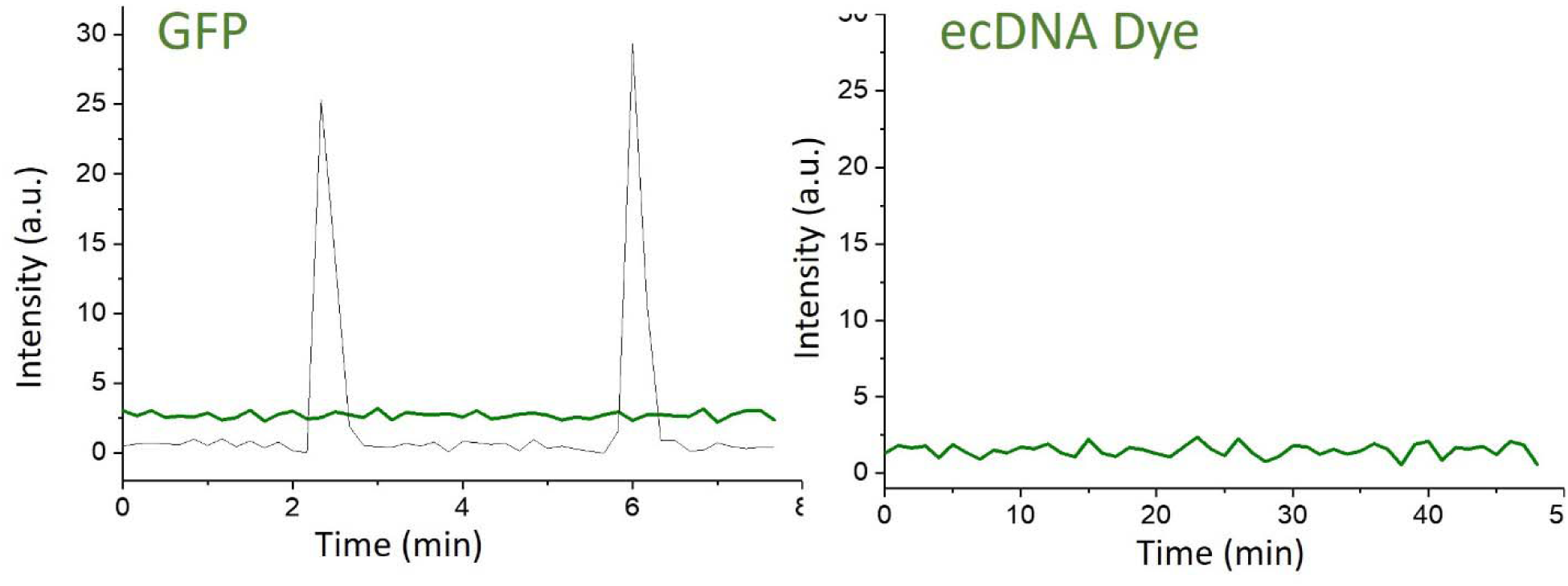
GFP labeling and extracellular DNA staining were used to verify the origin of the optical scattering, specifically to distinguish between surface versus bulk oscillations in the biofilm. Neither fluorescence imaging method revealed oscillatory signals. Simultaneously recorded gray signals, indicating the presence of original waves on the biofilm, were superimposed for comparison.

**Supplementary Figure 3.**
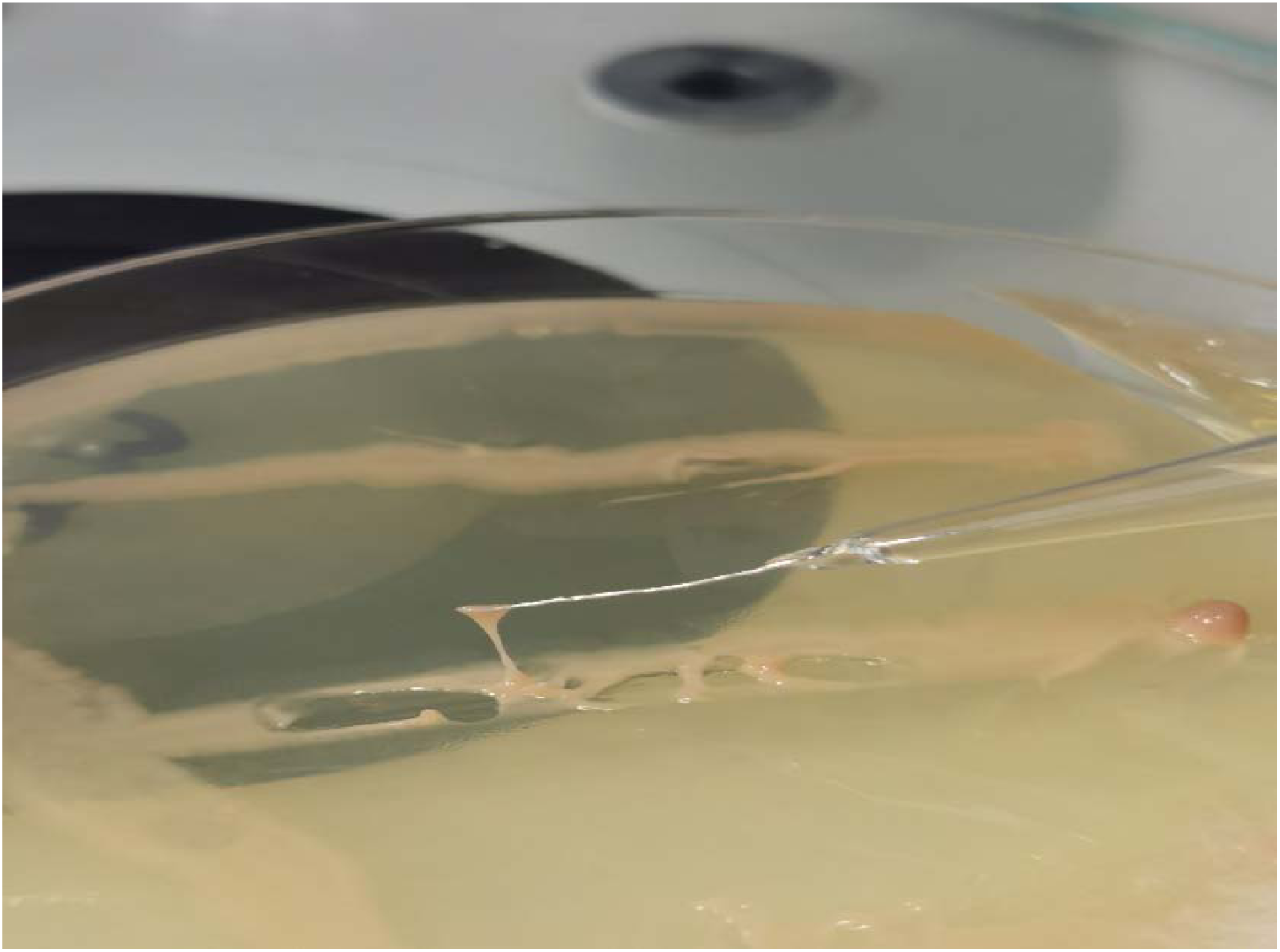
Image showing the elastic properties of the biofilm (PN) at the center of the colony. The biofilm was elastically stretched using a sharp needle. Unlike the finger-forming colony edge, this elastic central region generates spiral waves.

**Supplementary Figure 4.**
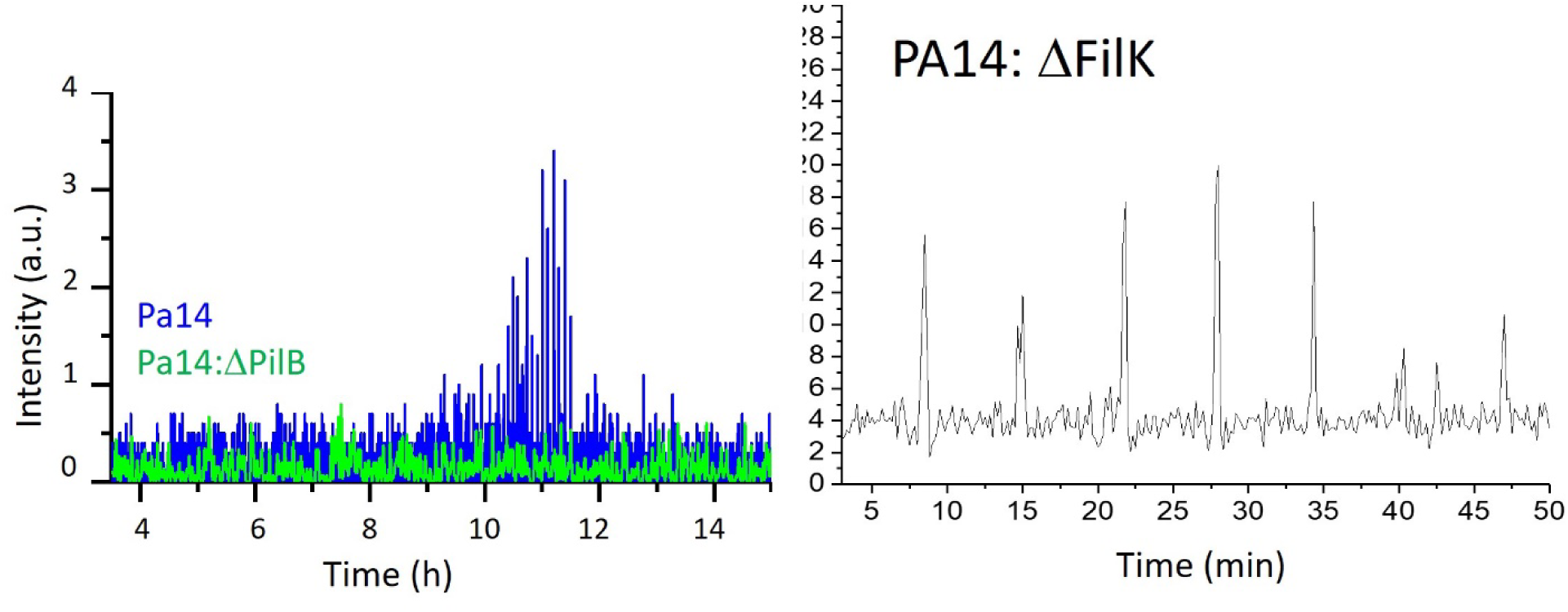
In PA14 strains, mutations in pilB and fliK were introduced to identify the processes responsible for driving the oscillations. The pilB mutation abolished wave formation, whereas wave emergence persisted in the fliK mutant, which lacks functional flagella.

**Supplementary Figure 5.**
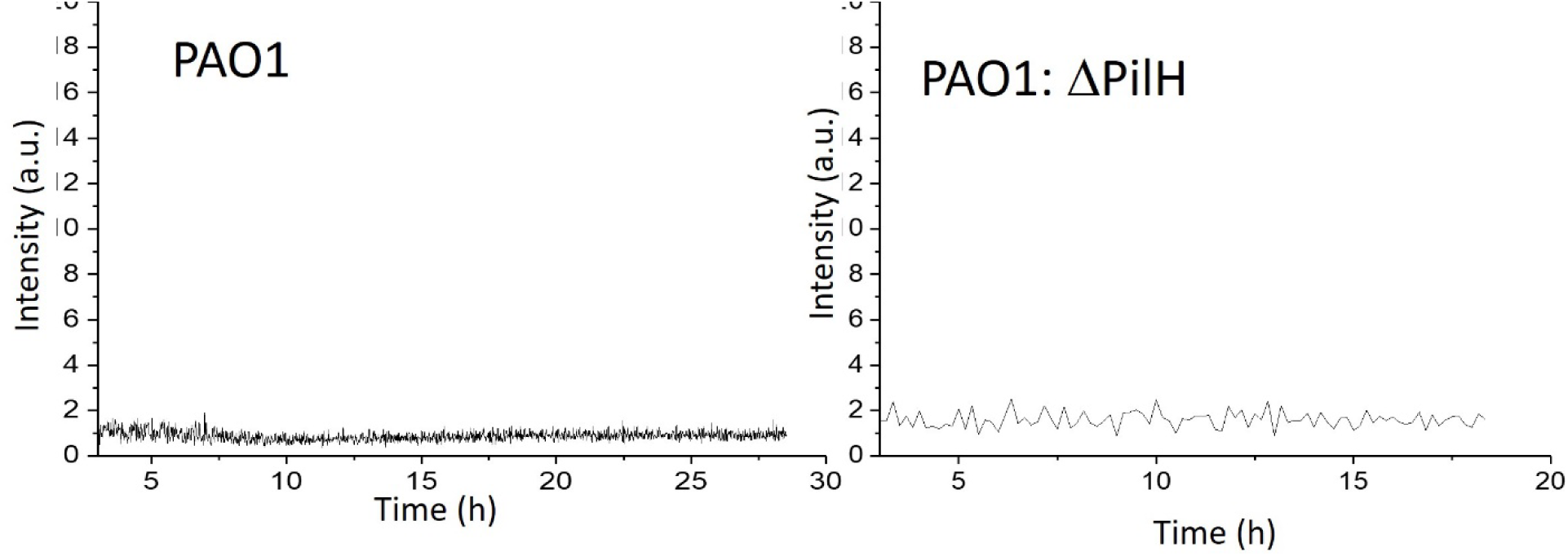
The PAO1 strain did not generate surface waves. Additionally, the hyperpiliated pilH mutant also failed to exhibit oscillatory wave behavior.

**Supplementary Figure 6.**
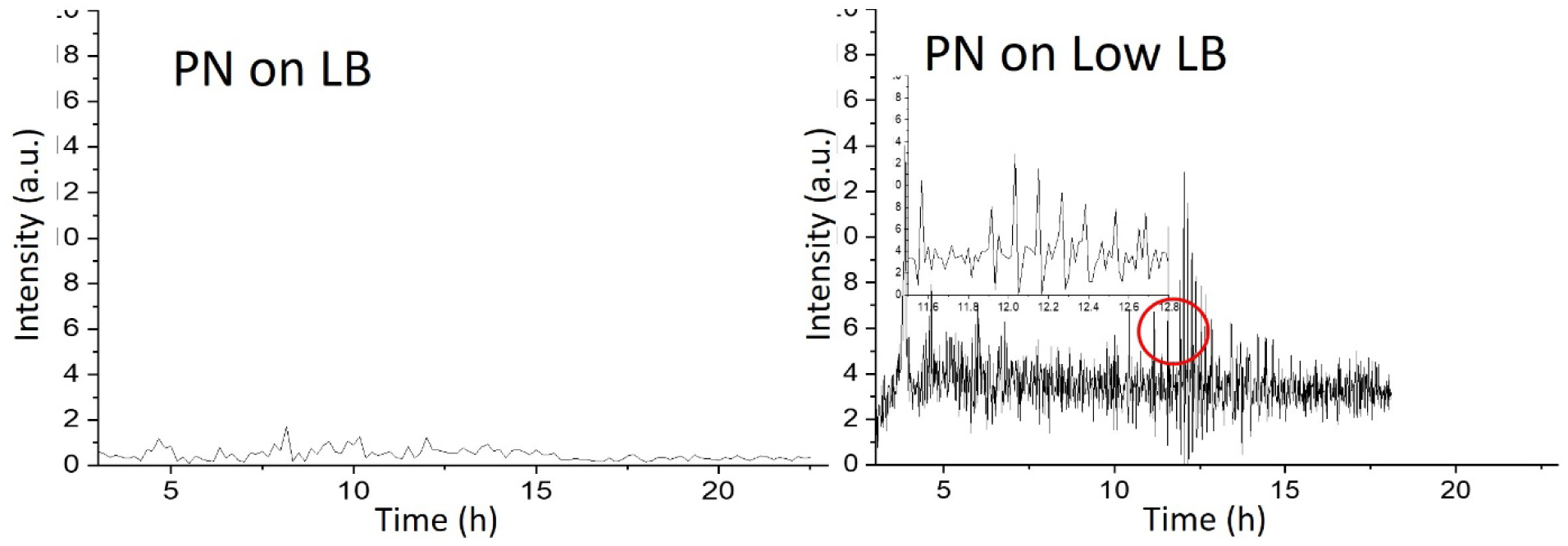
Regular LB plates do not provide sufficient conditions for wave generation in PN. However, eliminating yeast extract and replacing tryptone with a low concentration (0.2×) peptone restored wave formation.

**Supplementary Figure 7.**
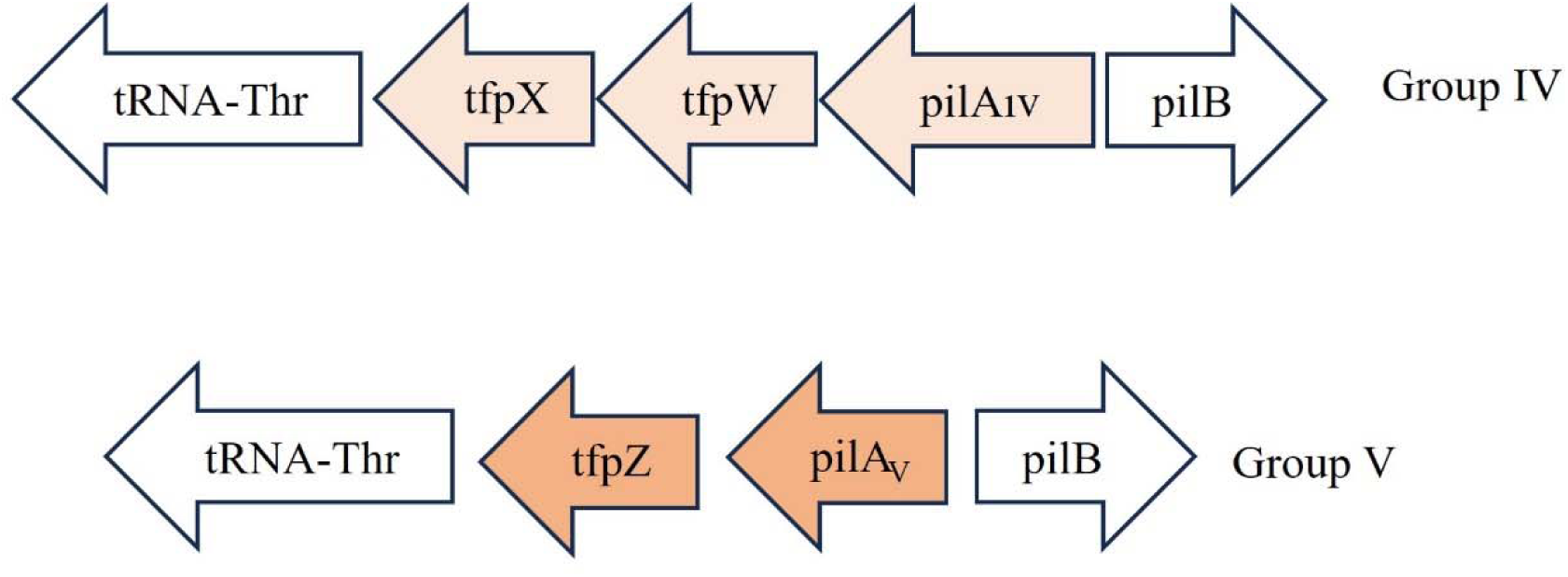
Comparative analysis of the genomic region encoding pili subunit groups reveals that PA14 contains Group 3 pili, while PN exhibits a Group 4 and 5-like structure. Both strains also encode accessory proteins associated with unknown pilus function.

**Supplementary Figure 8.**
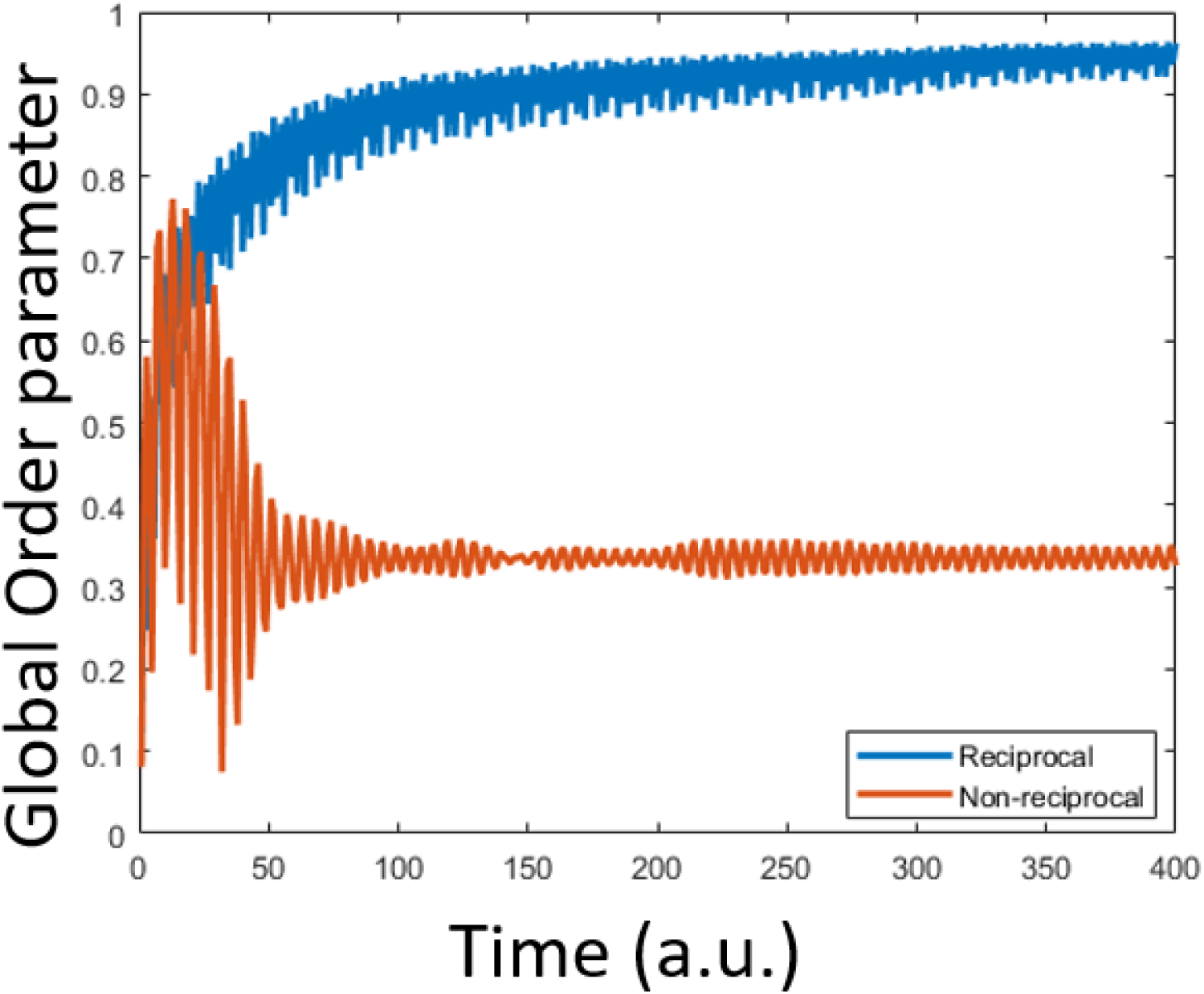
Global order parameter as a function of time for reciprocally and nonreciprocally coupled oscillators

**Supplementary Figure 9.**
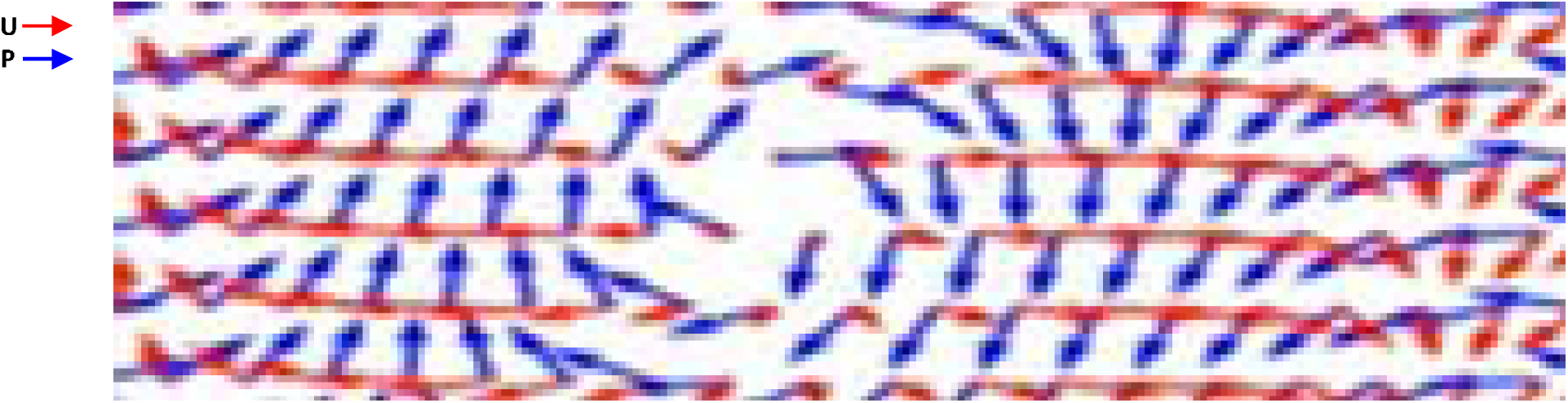
Vector configuration around a topological defect, where the displacement vector changes direction.

**Supplementary Figure 10.**
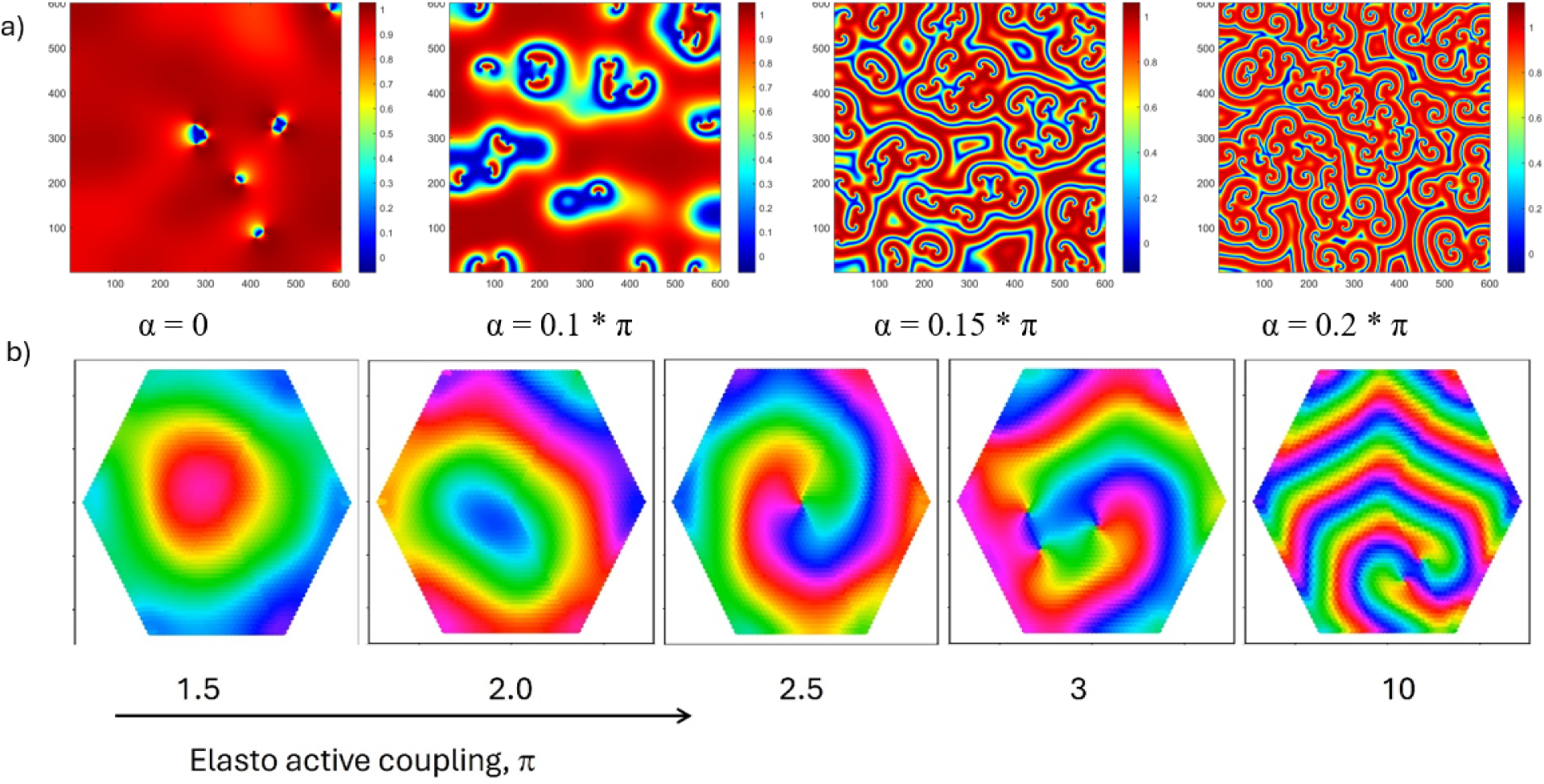
Defect dynamics as a function of the control parameter. (a) In the nonreciprocally coupled oscillator model, a large number of defects persist at high nonreciprocity, whereas at low nonreciprocity, oppositely charged (+/−) defect pairs annihilate. (b) In the active gel model, similarly, +/− defect pairs remain stable at high nonreciprocity, demonstrating consistent defect stabilization across both systems.

**Supplementary Figure 11.**
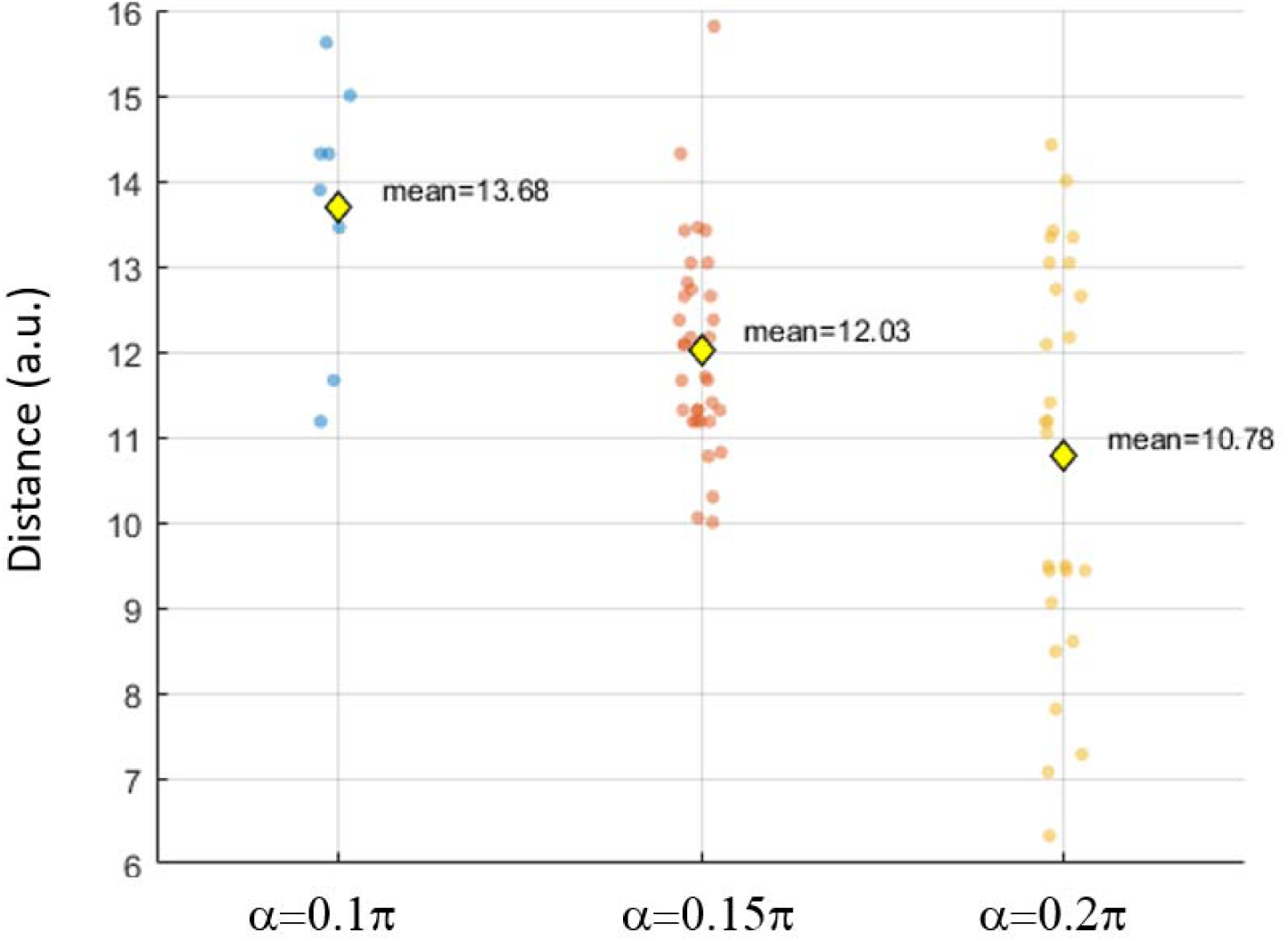
Separation between the defect pairs. The annihilation of +/− defect pairs, defect separation strongly depends on the nonreciprocity coefficient of the system. At high nonreciprocity, the system can sustain closely separated defect pairs. This regime corresponds to the dry condition of the biofilm.

**Supplementary Figure 12.**
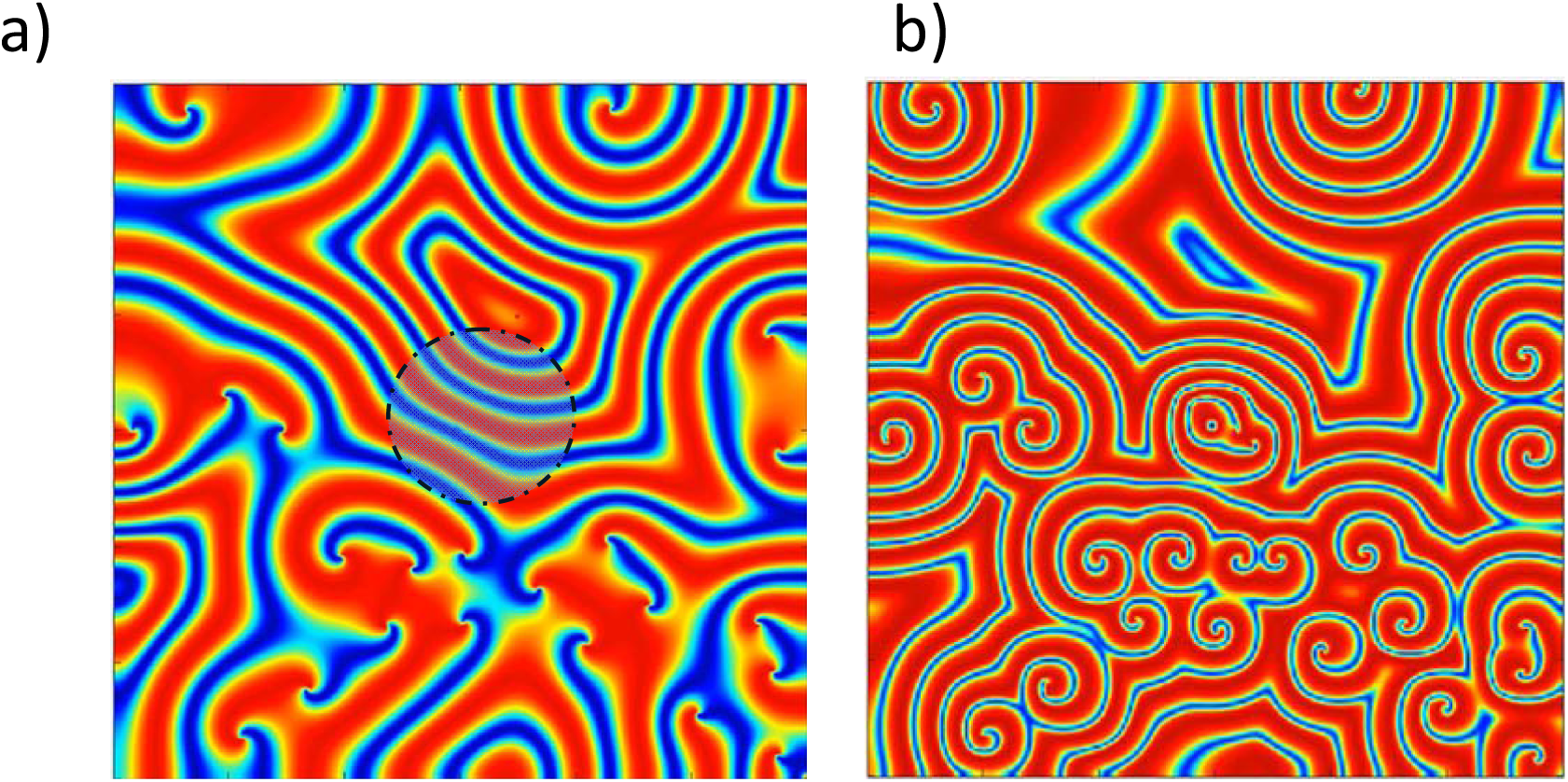
Simulation results of defect generation by introducing nonuniformities.

**Supplementary Figure 13.**
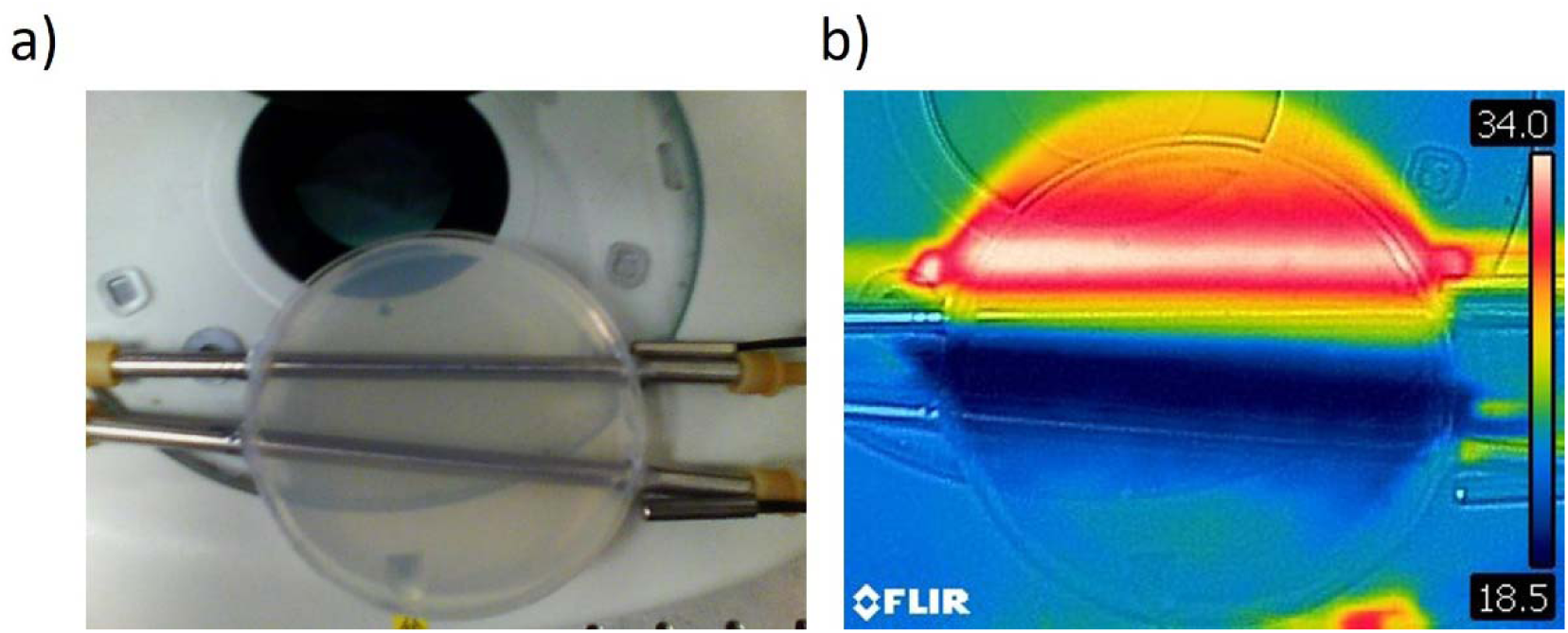
Optical and thermal images of an NGM plate equipped with metallic pipes. A closed-loop water pumping-heating system was used to control the local temperature and create a temperature gradient between the pipes. The spacing between the pipes was varied to identify the optimal gradient profile.

## Data availability

The critical experimental data generated or analyzed during this study are provided as supporting video files. We did not generate additional data sets.

## Software availability

All the codes used in the study are available online (https://github.com/akocabaslab/kuramoto-oscillators).

## Materials and Methods

### Analysis of Pilin Proteins and Accessory Genes

Genetic analyses were conducted using the complete genome sequence of Pseudomonas nitroreducens strain L4 chromosome^71^ (NCBI Reference Sequence: NZ_CP120376.1). The reference genomic region between pilB and tRNA-thr was specifically compared.

### Imaging System

Time-lapse imaging was performed using multiple microscopy techniques, selected for their capability to visualize surface topography effectively. We observed that oblique and polarized illumination techniques significantly enhanced the visibility of surface waves. During the evaporation process, the roughness of the biofilm significantly varies. During the evaporation process, the roughness of the biofilm varies significantly. We observed that oblique illumination is more effective on rough surfaces, whereas polarized imaging provides clearer results on flat surfaces. Oblique imaging was primarily conducted using Zeiss Smartzoom digital microscopes and Nikon Stereo SMZ18 microscopes equipped with contrast illumination. Additional polarized imaging utilized a Zeiss Axio Imager M2m. Successive images were captured at intervals of 10 seconds. Fluorescence imaging for GFP was conducted using 488 nm blue-light excitation with GFP emission filters, and DNA staining was visualized using RFP emission filters and imaged with an Andor EMCCD camera. The typical optical magnification was 10X.

### Image Processing

To enhance wave visibility, successive images taken at 10-second intervals were subtracted, and the resulting false-colored difference images were superimposed onto the original ones. The time-dependent responses of surface pulses were analyzed using ImageJ software.

### Bacterial growth and Preparation

The Pseudomonas nitroreducens (PN) bacterial strain used in this study was initially isolated from environmental samples and identified through 16S rRNA sequencing. Verification was performed through comparison with the corresponding strain obtained from DSMZ. For optical imaging, PN bacteria were grown overnight in LB broth at 21 °C on a shaker, and subsequently, 100 µl drops of bacterial suspension were placed onto 10-cm, 1-day-old, 1.4% solid agar, NGM plates. To ensure slow evaporation of excess moisture, humidity was maintained at approximately 80%, and plate lids were partially left open. Nematode growth medium (NGM) plates were prepared following the standard protocols.

### Numerical Simulation

Numerical simulations based on the Kuramoto model were performed using custom-developed MATLAB code with finite difference algorithms (https://github.com/akocabaslab/kuramoto-oscillators). The original code for the active solid model was obtained from prior literature and modified for improved visualization. Through extensive parameter exploration, we determined that large lattice structures with fixed boundary conditions effectively reproduced spiral and planar wave dynamics.

## Author contributions

B.A. and A.Ko. initiated the project and performed the experiments. B.A., Y.I.Y., and M.B. Y.G. developed and executed numerical modeling. B.K. conducted 16S DNA sequencing and GFP labeling of PN. N.G. performed genome sequence analysis. E.T.G. and A.Ü. developed and conducted imaging for the temperature gradient control system. S.K.Ö. and C.K. conceptualized non-Hermitian and PT-symmetry principles. A.Ko. and B.A. drafted the manuscript, with contributions from all authors toward the final manuscript.

## Acknowledgments

This work was supported by Airforce Office of Scientific Research (AFOSR) under Grant No: FA9550-22-1-0431 with Program Manager Ali Sayir and Tübitak Project No: 121C159. S.K.O acknowledges the support from AFOSR Multidisciplinary University Research Initiative (MURI) Award No. FA9550-21-1-0202. We thank Zeiss, Turkey division and Kenan Doğru for digital microscopy installation and training.

## Competing interests

The authors declare no competing interests.

## Video Captions

**Video 1.** Time-lapse imaging of active *Pseudomonas nitroreducens* biofilm surfaces generating metachronal waves. Global left-right symmetry is broken, and merging spiral waves at the edge drive unidirectional planar waves toward the biofilm center. The wave wavelength is approximately 100 µm. Associated with Figure 1a.

**Video 2.** Time-lapse imaging of circularly symmetric active biofilm surfaces generating inward-propagating metachronal waves. Associated with Figure 1b.

**Video 3.** High magnification (100X) time-lapse imaging of biofilm surfaces illustrating bacterial displacement and optical contrast changes.

**Video 4.** Time-lapse imaging of biofilm growth initiated from a single bacterium. After growth cessation, dense regions begin firing first, and waves dominate the biofilm surface.

**Video 5.** Time-lapse imaging of fingering formation at the biofilm edge. Bacteria remain motile and exhibit collective flow.

**Video 6.** Time-lapse imaging of the biofilm center generating propagating waves. Bacteria are attached to the surface; however, they periodically lift up and deform the surface, resembling Mexican-wave dynamics.

**Video 7.** Numerical simulation using the nonreciprocal Kuramoto model. Colors indicate oscillation phases. Spiral waves merge to form target waves, eventually converging into planar waves. Associated with Figure 2c.

**Video 8.** Numerical simulation based on the active solid model. Emergence of spiral and planar waves within the lattice. Associated with Figure 2e.

**Video 9** Time-lapse imaging of controlling the transition between waves to target and spiral by manipulating the moisture of biofilm surface. Associated with Figure 3d-g

**Video 10** Time-lapse imaging of a bacterial biofilm with broken left-right symmetry. Associated with Figure 6.

**Video 11.** Numerical simulation of propagating planar waves influenced by a frequency gradient. Associated with Figure 7c.

**Video 12.** Numerical simulation of defect dynamics triggered by nonuniformities at low nonreciprocal conditions (α=0.1π).

**Video 13.** Numerical simulation of defect generation triggered by nonuniformities at high (α=0.2π) nonreciprocal conditions.

**Video 14.** Time-lapse imaging of controlling propagating planar waves under a varying temperature gradient. Associated with Figure 7.

## Competing interests

Authors declare no competing interests.

**Table 1:** List of strains used in this study.

| Strain | Parent | Operation | Genotype |
| --- | --- | --- | --- |
| BAK132 | PN | Isolated from environmental samples, transformed pUCP18-MCSgfpmut3 | *ampR* |
| BAK133 | PA14 | Received from F. M. Ausubel lab |  |
| BAK134 | LD2222 | Received from Lars Dietrich Lab<sup>72</sup> | PA14 ΔpilB |
| BAK135 | LD2221 | Received from Lars Dietrich Lab<sup>72</sup> | PA14 ΔflgK |
| BAK136 | PAO1 PilH | Received from Joanne Engel | ΔPilH |
| BAK | PAO1 | Received from DSMZ 10145GFP |  |

## Notes

### Competing Interest Statement

The authors have declared no competing interest.

### Summary of Updates

The new findings of the manuscript are supported by additional experimental and numerical results.

